# Free fatty-acid receptor 4 inhibitory signaling in delta cells regulates islet hormone secretion in mice

**DOI:** 10.1101/2020.07.17.208637

**Authors:** Marine L. Croze, Marcus F. Flisher, Arthur Guillaume, Caroline Tremblay, Glyn M. Noguchi, Sabrina Granziera, Kevin Vivot, Vincent C. Castillo, Scott A. Campbell, Julien Ghislain, Mark O. Huising, Vincent Poitout

**Author notes:** Corresponding author: Vincent Poitout, DVM, PhD, CRCHUM, 900 rue St Denis, Montréal, QC, H2X 0A9 - CANADA.

## Abstract

**Objective:** Maintenance of glucose homeostasis requires the precise regulation of hormone secretion from the endocrine pancreas. Free fatty-acid receptor 4 (FFAR4/GPR120) is a G protein-coupled receptor whose activation in islets of Langerhans promotes insulin and glucagon secretion and inhibits somatostatin secretion. However, the contribution of individual islet cell types (α, β, and δ cells) to the insulinotropic and glucagonotropic effects of GPR120 remains unclear. As *gpr120* mRNA is enriched in somatostatin-secreting δ cells, we hypothesized that GPR120 activation stimulates insulin and glucagon secretion via inhibition of somatostatin release.

**Methods:** Glucose tolerance tests were performed in mice after administration of the selective GPR120 agonist Compound A. Insulin, glucagon and somatostatin secretion were measured in static incubations of isolated mouse islets in response to endogenous (ω-3 polyunsaturated fatty acids) and/or pharmacological (Compound A and AZ-13581837) GPR120 agonists. The effect of Compound A on hormone secretion was tested further in islets isolated from mice with global or somatostatin cell-specific knockout of *gpr120*. *Gpr120* expression was assessed in pancreatic sections by RNA in situ hybridization. Cyclic AMP (cAMP) and calcium dynamics in response to pharmacological GPR120 agonists were measured specifically in α, β and δ cells in intact islets using cAMPER and GCaMP6 reporter mice, respectively.

**Results:** Acute exposure to Compound A increased glucose tolerance and circulating insulin and glucagon levels in vivo. Endogenous and/or pharmacological and GPR120 agonists reduced somatostatin secretion in isolated islets and concomitantly demonstrated dose-dependent potentiation of glucose-stimulated insulin secretion and arginine-stimulated glucagon secretion. *Gpr120* was enriched in δ cells. Pharmacological GPR120 agonists reduced cAMP and calcium levels in δ cells but increased these signals in α and β cells. Compound A-mediated inhibition of somatostatin secretion was insensitive to pertussis toxin. The effect of Compound A on hormone secretion was completely absent in islets from mice with either global or somatostatin cell-specific deletion of *gpr120* and was partially reduced upon blockade of somatostatin receptor signaling by cyclosomatostatin.

**Conclusions:** Inhibitory GPR120 signaling in δ cells contributes to both insulin and glucagon secretion in part via mitigating somatostatin release.

## 1. INTRODUCTION

G protein-coupled receptors are validated targets for the treatment of type 2 diabetes [1] and among these, the long-chain FA^1^ receptor GPR120/FFAR4 has been the subject of increasing interest in recent years as its activation has numerous beneficial effects on glucose and energy homeostasis in preclinical models [2]. In rodents, GPR120 activation alleviates obesity-induced chronic inflammation and associated insulin resistance [3; 4], promotes adipogenesis [5–7] and brown adipose tissue thermogenesis [8; 9], inhibits lipolysis in white adipose tissue [10], regulates food intake [11], and modulates enteroendocrine hormone secretion, including ghrelin [12–14], glucagon-like peptide-1 [15], glucose-dependent insulinotropic polypeptide [16], cholecystokinin [17; 18] and SST [19].

GPR120 is also reportedly expressed in islet α, β, δ and γ cells where its activation mitigates β-cell dysfunction [20] and apoptosis [21] and modulates islet hormone secretion. GPR120 activation promotes GSIS [22–26]; potentiates glucagon secretion [26; 27]; inhibits GSSS [28]; and stimulates PP secretion [29].

GPR120 signaling was reported to promote insulin secretion in insulin-secreting cell lines via intracellular calcium mobilization [22; 24]. Yet, transcriptomic profiling and RT-PCR indicate that *gpr120* is primarily expressed in the δ cell with lower levels detected in α and β cells [21; 29–32]. Preferential expression of *gpr120* in islet δ cells was confirmed by knock-in of the LacZ reporter into the *gpr120* locus in mice [28]. However, the functional contribution of GPR120 signaling in individual islet endocrine cell types to the net effect of its activation on insulin and glucagon secretion remains unknown. To clarify the role of GPR120 in islet hormone secretion and define the precise contribution of δ-cell GPR120 signaling in these processes, we measured insulin, glucagon and SST secretion in response to natural and synthetic GPR120 agonists in isolated islets from WT, whole-body *gpr120* KO, and SST-cell specific *gpr120* KO mice; determined the cellular localization of *gpr120* in pancreatic sections; investigated calcium fluxes and cAMP generation in response to GPR120 agonists in α, β and δ cells directly within intact islets; and assessed the contributions of PTX-sensitive G proteins and SST.

## 2. MATERIALS AND METHODS

### 2.1 Reagents and solutions

RPMI-1640 and FBS were from Life Technologies Inc. (Burlington, ON, Canada). Penicillin/Streptomycin was from Multicell Wisent Inc (Saint-Jean-Baptiste, QC, Canada). FA-free BSA was from Equitech-Bio (Kerrville, TX, USA). Cpd A was from Cayman Chemical (Ann Arbor, MI, USA). AZ was generously provided by AstraZeneca (Gothenburg, Sweden). Insulin and glucagon RIA kits were from MilliporeSigma (Billerica, MA, USA). SST RIA kits were from Eurodiagnostica (Malmö, Sweden). Insulin and glucagon ELISA kits were from Alpco Diagnostics (Salem, NH, USA) and Mercodia (Uppsala, Sweden), respectively. PTX was from List labs (Campbell, CA, USA). cSST was from Tocris bioscience (Minneapolis, MN, USA). Aprotinin was from Roche Diagnostics (Rotkreuz, Switzerland). All other reagents were from MilliporeSigma unless otherwise specified.

### 2.2 Animals

All procedures involving animals were approved by the Institutional Committee for the Protection of Animals (IACUC) at the Centre Hospitalier de l’Université de Montréal, with the exception of GCaMP6 or CAMPer reporter mice, which were maintained for islet collection under the supervision of the IACUC of UC Davis. Animals at each institution were handled in accordance with the National Institutes of Health guide for the care and use of Laboratory animals. All mice were housed under controlled temperature on a 12h light/dark cycle with unrestricted access to water and standard laboratory chow and were sacrificed at 10-12 weeks of age for islet isolation. C57BL/6N male mice were purchased from Charles River (Saint-Constant, QC, Canada).

#### Whole-body gpr120 KO mice

Mice carrying LoxP sites flanking exon 1 of *gpr120* (Gpr120^floxNeofrt^) were obtained from Ingenious Targeting Laboratory, Ronkonkoma, NY, U.S.A. The neo cassette was removed by crossing the mice to ROSA26:FLPe mice (129S4/SvJaeSor-Gt(ROSA)26Sortm1(FLP1)Dym/J, The Jackson Laboratory, Bar Harbor, ME, USA), and the resulting mice were back-crossed onto a C57BL/6N background for more than 9 generations. Unexpectedly, homozygous Gpr120^flox^ (prev-Flox) mice displayed an important reduction in *gpr120* gene expression and function in islets (**Supplementary Fig. 1A & B**) and other organs (data not shown), suggesting abnormal transcription of *gpr120* resulting from insertion of the LoxP sites and making them unsuitable for conditional KO studies. These Gpr120^flox^ mice were crossed with transgenic E2A-Cre mice (B6.FVB-Tg(EIIa-cre)C5379Lmgd/J, The Jackson Laboratory) to remove exon 1 (Gpr120^Δ^) to generate whole-body *gpr120* KO animals (Gpr120KO). As the E2A-Cre mouse was of a mixed B/6N and B/6J background carrying the NNT mutation (Nnt^C57BL/6J^), only Gpr120^Δ^ mice lacking E2A-Cre and Nnt^C57BL/6J^ were kept for subsequent crossings. Male WT and Gpr120KO experimental animals were generated by crossing heterozygous Gpr120^Δ^ mice. Genotyping primers are listed in **Supplementary Table 1**. Animals were born at the expected Mendelian ratio and expression of *gpr120* was completely eliminated in KO islets (**Supplementary Fig. 1C**).

#### Whole-body gpr40 KO mice

Male WT and *gpr40* KO mice were generated and genotyped as previously described [33].

#### SST cell-specific gpr120 KO mice

Mice carrying LoxP sites flanking exon 1 and approximately 1.5 kb of sequence upstream of exon 1 (promoter region) of *Ffar4* on a C57BL/6N background (C57BL/6-Ffar4^tm1.1Mrl^) were purchased from Taconic Biosciences (hereafter designated Gpr120^+/fl^ or Gpr120^fl/fl^) and crossed with heterozygous SST-Cre mice (B6N.Cg-Sst^tm2.1(cre)Zjh^/J, Jackson Laboratory) also on the C57BL/6N background. Gpr120^+/fl^ and SST-Cre;Gpr120^+/fl^ mice were crossed to generate experimental groups. Male WT, Gpr120^fl/fl^ (Flox), SST-Cre (Cre) and SST-Cre;Gpr120^fl/fl^ mice were used for secretion experiments. Since experiments were performed in isolated islets in which only δ cells express SST, SST-Cre;Gpr120^fl/fl^ mice are thereafter referred to as δGpr120KO mice. Genotyping primers are listed in **Supplementary Table 1**. Animals were born at the expected Mendelian ratio. Females of the 4 genotypes were used for qPCR experiments. *Gpr120* mRNA levels were significantly reduced in δGpr120KO islets, in accordance with the predominant expression of *gpr120* in δ cells (**Supplementary Fig. 1D**). *Gpr120* mRNA levels were similar in islets from WT, Gpr120Flox and SST-Cre mice.

#### cAMPER and GCaMP6 reporter mice

We used the following Cre drivers to express fluorescent biosensors for calcium (GCaMP6s; Gt(ROSA)26Sor^tm96(CAG-GCaMP6s)Hze^) [34] or cAMP (C57BL/6-Gt(ROSA)26Sor^tm1(CAG-ECFP*/Rapgef3/Venus*)Kama^/J) (cAMPER, The Jackson Laboratory, strain #032205) [35] directly and specifically in α cells (B6;129S4-Gcg^em1(cre/ERT2)Khk^/Mmjax) [36], β cells (B6.FVB(Cg)-Tg(Ucn3-cre)KF43Gsat/Mmucd) [37], or δ cells (B6N.Cg-Sst^tm2.1(cre)Zjh^/J).

### 2.3 Fluorescence mRNA in situ hybridization

mRNAs were visualized by fluorescence in situ hybridization using the RNAscope Fluorescent Multiplex Kit (Advanced Cell Diagnostics, Inc., Hayward, California, USA) on fixed and frozen 12-week-old male C57Bl/6N pancreatic cryosections. Briefly, pancreata were fixed overnight in 4% paraformaldehyde and cryoprotected overnight in 30% sucrose. Tissues were then embedded in OCT, frozen, sectioned at 8 μm, and mounted on Superfrost Plus slides (Life Technologies, Carlsbad, CA, USA). The following RNAscope probes were used: Mm-Ffar4-O1 (Cat No. 447041), Mm-Ins2-O1-C2 (Cat No. 497811-C2), Mm-Sst-C4 (Cat No. 404631-C4) and Mm-Gcg-C3 (Cat No. 400601-C3). Hybridization and fluorescent detection were performed according to the manufacturer’s instructions. 20X and 63X images were acquired with an inverted confocal microscope (Leica Microsystem, Mannheim, Germany).

### 2.4 Islet isolation

Mouse islets were isolated by collagenase digestion and dextran density gradient centrifugation as described previously [33] and allowed to recover overnight in RPMI 1640 supplemented with 10% (wt/vol) FBS, 100 U/ml penicillin/streptomycin and 11 mM glucose.

### 2.5 RNA extraction and quantitative RT-PCR

Total RNA was extracted from batches of 120-200 islets using the RNeasy micro kit (Qiagen, Valencia, CA). RNA was quantified by spectrophotometry using a NanoDrop 2000 (Life Technologies) and 0.4-1.0 μg of RNA was reverse transcribed. Real-time PCR was performed by using QuantiTect SYBR Green PCR kit (Qiagen, Hilden, Germany). Results were normalized to *cyclophilin A* (*ppia*) mRNA levels and normalized to the levels in control islets. Primer sequences are listed in **Supplementary Table 2**.

### 2.6 Static incubations for insulin and SST secretion

After overnight recovery, islets were incubated in KRBH (pH 7.4) with 0.1% (w/v) FA-free BSA and 2.8 mM glucose for 20 min. Triplicate batches of 20 islets each were then incubated for an additional 20 min in KRBH, 0.1% FA-free BSA, 2.8 mM glucose, followed by a 1-h static incubation in KRBH in the presence of 2.8 or 16.7 mM glucose, with or without synthetic GPR120 agonists (10, 20 or 50 μM Cpd A; 0.1, 1, 5 or 10 μM AZ) or endogenous ligands (ALA, EPA, DHA, 0.1 mM), as indicated in the figure legends. We selected Cpd A and AZ among the different GPR120 agonists because of their selectivity towards GPR120 versus GPR40 [4; 23]. Islets were exposed to PTX (100 ng/ml) during the overnight recovery for 16 h as indicated in **Fig. 8**. cSST (10 μM) was included during the last 20-min preincubation and 1-h static incubation as indicated in **Fig. 8**. Secreted SST and insulin were measured in the supernatant by RIA. Intracellular insulin content was measured after acid–alcohol extraction.

### 2.7 Static incubations for glucagon and SST secretion

After overnight recovery, islets were incubated in KRBH (pH 7.4) with 0.1% (w/v) FA-free BSA and 5.5 mM glucose for 20 min. Triplicate batches of 20 islets each were then incubated an additional 20 min in KRBH, 0.1% FA-free BSA, 5.5 mM glucose, followed by a 1-h static incubation in KRBH in the presence of 1 mM glucose, with or without 10 mM L-arginine, GPR120 agonists (10 or 50 μM Cpd A,; 0.1, 1, 5 or 10 μM AZ). cSST (10 μM) was included during the last 20-min preincubation and 1-h static incubation as indicated in **Fig. 8**. Secreted SST and glucagon were measured in the supernatant by RIA. Intracellular glucagon content was measured after acid–alcohol extraction.

### 2.8 Calcium and cAMP signaling in α, β, and δ cells

As previously described [37], following isolation islets were cultured overnight after which islets were placed in 35mm glass-bottom dishes (#1.5; MatTek Corporation, Ashland, MA, USA), allowed to attach overnight and imaged in x, y, z and t on a Nikon A1R+ confocal microscope using a 40X or 60X lens with a long working distance under continuous perfusion. For calcium imaging, islets were excited by a 488 nm excitation line, with the emitted signal collected through a 525/50 nm BP filter, which each protocol concluding with a 30 mM KCl pulse to demonstrate viability and responsiveness throughout the treatment. Individual cells in individual z-planes were defined as regions of interest (ROI) and the green fluorescence intensity within the ROIs was plotted over time as a measure of calcium activity. To trace cAMP islets were processed and imaged essentially as described above for calcium tracing, but exciting with a 445 laser line while simultaneously detecting CFP (485/40 nm BP) and YFP (525/50 nm BP) emission with two parallel detectors. Forskolin was used instead of KCl as our positive indicator of cell viability and ability to mount a cAMP response.

### 2.9 Metabolic tests

Cpd A (60 mg/kg BW) was administered orally in Cremophor-EtOH-water (1/1/18, v/v/v) to 4-h fasted mice 30 min prior to oral glucose administration (1 g/kg BW) or immediately before intraperitoneal L-arginine injection (1.25 g/kg BW). Tail blood glucose was measured using a hand-held Accu-Chek glucometer (Roche, Indianapolis, IN). For glucagon measurements, aprotinin (0.5 KIU/μL) was added immediately after collection and plasma samples were immediately snap frozen in liquid nitrogen. Plasma insulin and glucagon were measured by ELISA.

### 2.10 Statistical analyses

Data are expressed as mean ± SEM. Significance was tested using standard one-way ANOVA, Brown-Forsythe and Welch ANOVA and corrections, in cases of variance heterogeneity, or two-way ANOVA with post hoc adjustment for multiple comparisons, as appropriate, using GraphPad Instat (GraphPad Software, San Diego, CA). Tukey or Dunnett post hoc tests were performed as indicated in figure legends. P<0.05 was considered significant.

## 3. RESULTS

### 3.1 Cpd A acutely improves glucose tolerance and potentiates insulin and glucagon secretion in vivo

To assess the effect of GPR120 activation on insulin secretion in vivo, Cpd A was administered orally to C57BL/6N mice 30 min prior to an oral glucose or immediately before an arginine tolerance test. Cpd A-treated mice displayed significantly lower glycemia 30 min after Cpd A administration and at all time points thereafter (**Fig. 1A & B**). The improved glucose tolerance of Cpd A-treated mice was associated with increased insulin levels at t = 30 min (**Fig. 1C & D**). Likewise, Cpd A potentiated arginine-induced glucagon secretion (**Fig. 1E**). Surprisingly, a corresponding increase in blood glucose was not detected, instead Cpd A-treated mice had significantly lower blood glucose 30 min after arginine administration (**Fig. 1F**). A Cpd A-dependent increase in insulin levels during the test may account for the decrease in blood glucose levels at 30 min. These results show that Cpd A improves glucose tolerance and increases insulin and glucagon secretion in mice.

**Figure 1.**
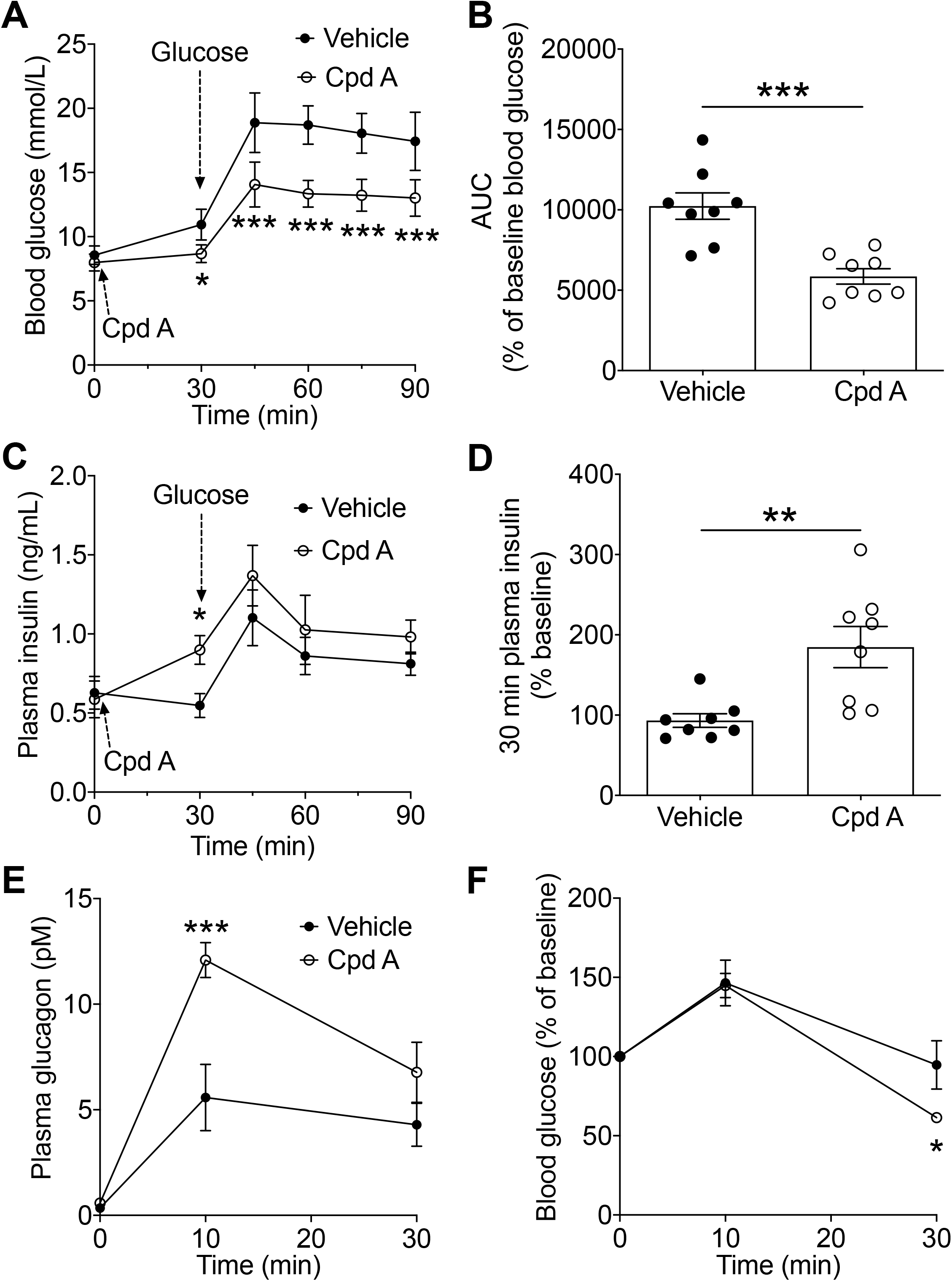
Cpd A potentiates insulin secretion and arginine-induced glucagon secretion in mice. Blood glucose (A), area under the curve (AUC) of blood glucose (B) and plasma insulin (C and D) in mice during an oral glucose tolerance test (1g/kg) performed 30 min after oral administration of Cpd A (60 mg/kg) or vehicle (Cremophor-EtOH-water). Data are mean +/− SEM of 8 animals per group. Plasma glucagon (E) and blood glucose (F) in C57BL/6N mice during an arginine test (1.25 g/kg, ip) performed after oral administration of Cpd A (60 mg/kg) or vehicle. Data are expressed as mean +/− SEM of 7 animals per group. * P<0.05, ** P<0.005 and *** P<0.0005 compared to vehicle group following two-way ANOVA with Tukey’s post hoc adjustment for multiple comparisons.

### 3.2 GPR120 activation potentiates GSIS and inhibits GSSS in isolated mouse islets

To assess whether intra-islet GPR120 activation controls islet hormone secretion we measured insulin and SST secretion in isolated WT mouse islets in response to glucose alone or in the presence of synthetic (Cpd A or AZ) or naturally-occuring ω-3 LCFA (ALA, EPA or DHA) GPR120 agonists. High glucose (16.7mM) significantly increased both insulin (**Fig. 2A & Supplementary Fig. 2A**) and SST (**Fig. 2B**) secretion compared to the low glucose (2.8 mM) condition, as expected. Both Cpd A (**Fig. 2A & B**) and AZ (**Fig. 2C & D**) potentiated GSIS (**Fig. 2A & C & Supplementary Fig. 2A & C**) and simultaneously inhibited GSSS (**Fig. 2B & D**). Interestingly, inhibition of SST secretion by both Cpd A and AZ was already maximal at the lowest concentrations used (**Fig. 2B & D**) whereas stimulation of insulin secretion was dose-dependent (**Fig. 2. A & C**), suggesting the contribution of SST-independent mechanisms to the regulation of insulin release. Endogenous GPR120 agonists (ALA, EPA, DHA) also potentiated GSIS (**Fig. 2E**) and inhibited GSSS (**Fig. 2F**). Although endogenous agonists were less efficient than the synthetic agonists in reducing GSSS, they displayed a greater potentiation of GSIS which is likely due to the combined activation of both GPR120 and GPR40 signaling pathways and non-receptor mediated effects due to intracellular metabolism. Cpd A, AZ and ω-3 FAs did not affect insulin content at any of the concentrations tested (**Supplementary Fig. 2B, D & F**).

**Figure 2.**
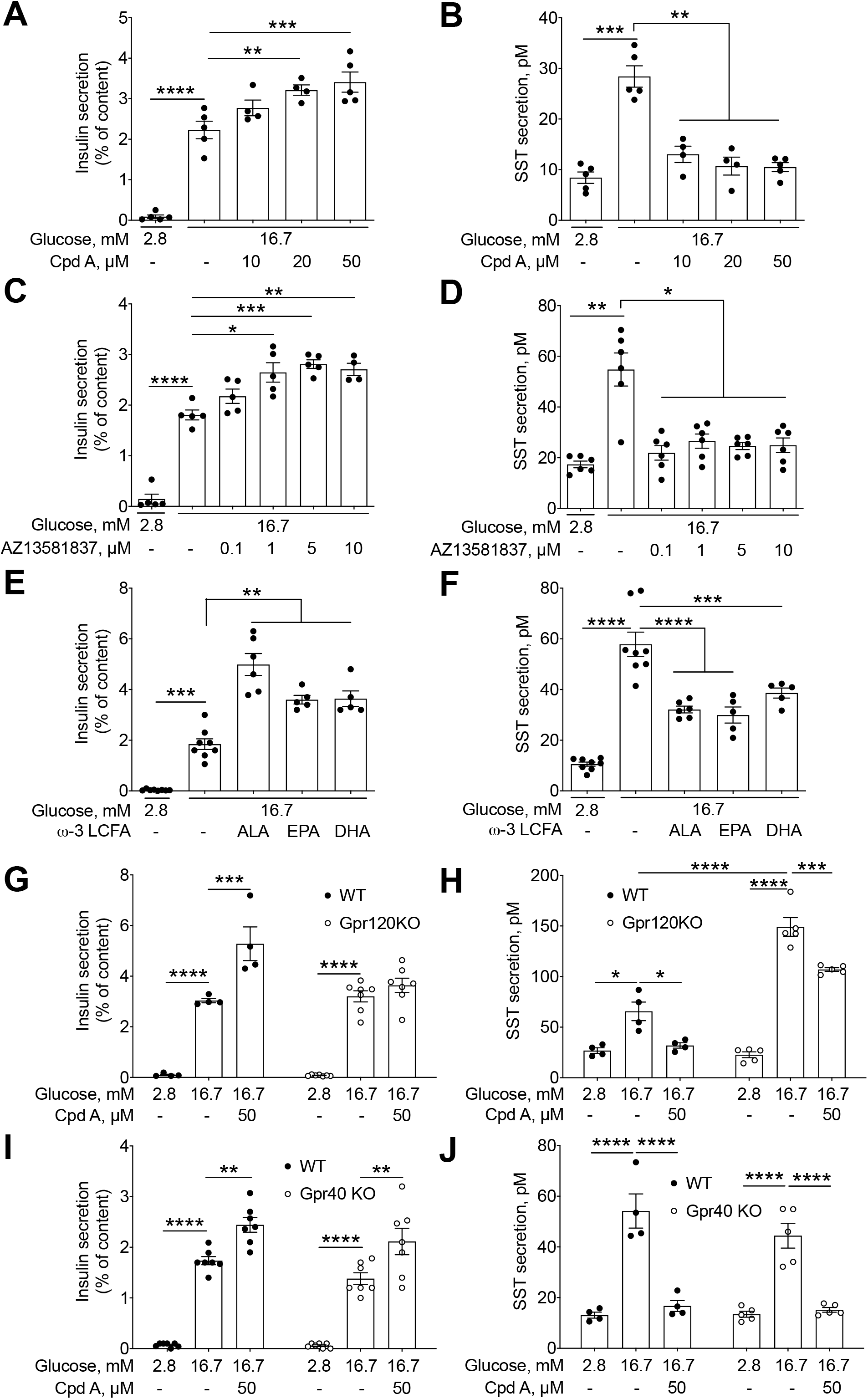
GPR120 activation potentiates GSIS and inhibits GSSS. Insulin secretion presented as a percentage of islet insulin content (A, C, E) and SST secretion measured in parallel on the same isolated islets (B, D, F) were assessed in 1-h static incubations in response to 2.8 or 16.7 mM glucose with or without the GPR120 agonists Cpd A (10-50 μM) (A, B), AZ (0.1-10 μM) (C, D) or the ω-3 fatty acids, alpha-linolenic acid (ALA), eicosapentaenoic acid (EPA) and docosahexaenoic acid (DHA) (100 μM) (E, F). Insulin and SST secretion in response to 2.8 or 16.7 mM glucose with or without Cpd A (50 μM) was measured in isolated islets from Gpr120KO (G, H) and Gpr40KO (I, J) mice and WT littermate controls. Data represent individual values and are expressed as mean +/− SEM of 4-8 independent experiments. * P<0.05, ** P<0.005, ***P<0.0005 or ****P<0.0001 between groups following one-way ANOVA (A-F) or two-way ANOVA (G-J) with Dunnett’s (A-F, versus 16.7-EtOH condition) or Tukey’s (G-J) post hoc adjustment for multiple comparisons and Welch/Brown-Forsythe correction when necessary to compensate for SD variances.

To test the selectivity of Cpd A towards GPR120, we treated Gpr120KO islets with the highest concentration of Cpd A (50 μM). The potentiation of GSIS induced by Cpd A in WT islets was absent in Gpr120KO islets (**Fig. 2G & Supplementary Fig. 2G**) without affecting insulin content (**Supplementary Fig. 2H**), confirming the requirement for GPR120 in the insulinotropic effect of Cpd A. Likewise, the inhibitory effect of Cpd A on GSSS was largely abrogated in Gpr120KO islets (**Fig. 2H**). Residual Cpd A-mediated inhibition of SST secretion in Gpr120KO islets suggests mild off-target effects of Cpd A in these experiments. Interestingly, GSSS was significantly higher in Gpr120KO versus WT islets with no difference in basal (2.8 mM glucose) SST secretion (**Fig. 2H**). This might be explained by the paracrine/autocrine action of islet-derived GPR120 ligands that reduce SST secretion in WT but not Gpr120KO islets.

In addition to their involvement in GSIS regulation, GPR120 and GPR40 are activated by an overlapping set of ligands [25]. Consequently, most GPR120 agonists activate GPR40 at high concentrations. To verify that GPR40 signaling does not contribute to the effect of Cpd A on GSIS, we measured insulin secretion in response to 50 μM Cpd A in Gpr40KO islets. The effects of Cpd A on GSIS (**Fig. 2I & Supplementary Fig. 2I**), GSSS (**Fig. 2J**) and insulin content (**Supplementary Fig. 2J**) were similar in WT and Gpr40KO islets, ruling out a contribution of GPR40 signaling.

### 3.3 GPR120 activation potentiates arginine-stimulated glucagon secretion and inhibits SST secretion in low glucose conditions in isolated mouse islets

Glucagon secretion was stimulated by 10 mM arginine in the presence of 1 mM glucose, and Cpd A dose-dependently potentiated arginine-induced glucagon secretion (**Fig. 3A & Supplementary Fig. 3A**) while inhibiting SST secretion (**Fig. 3B**). Similarly, AZ dose-dependently potentiated arginine-induced glucagon secretion (**Fig. 3C & Supplementary Fig. 3C**). Neither Cpd A nor AZ affected glucagon content at any of the concentrations tested (**Supplementary Fig. 3B & D**). Arginine-induced glucagon secretion was significantly decreased in Gpr120KO but not Gpr40KO islets (**Fig. 3D & F & Supplementary Fig. 3E & G**). Gpr120KO (**Fig. 3E)**, but not Gpr40KO (**Fig. 3G**) islets also exhibit higher SST secretion in response to arginine. These results are consistent with the presence of islet-derived GPR120 agonists that promote glucagon and inhibit SST secretion in WT and Gpr40KO but not Gpr120KO islets. As expected, the glucagonotropic effect of Cpd A was lost in Gpr120KO (**Fig. 3D & Supplementary Fig. 3E**) but maintained in Gpr40KO (**Fig. 3F & Supplementary Fig. 3G**) islets without effects on glucagon content (**Supplementary Fig. 3F & H**). Hence, GPR120 but not GPR40 is required for the potentiation of glucagon secretion by Cpd A.

**Figure 3.**
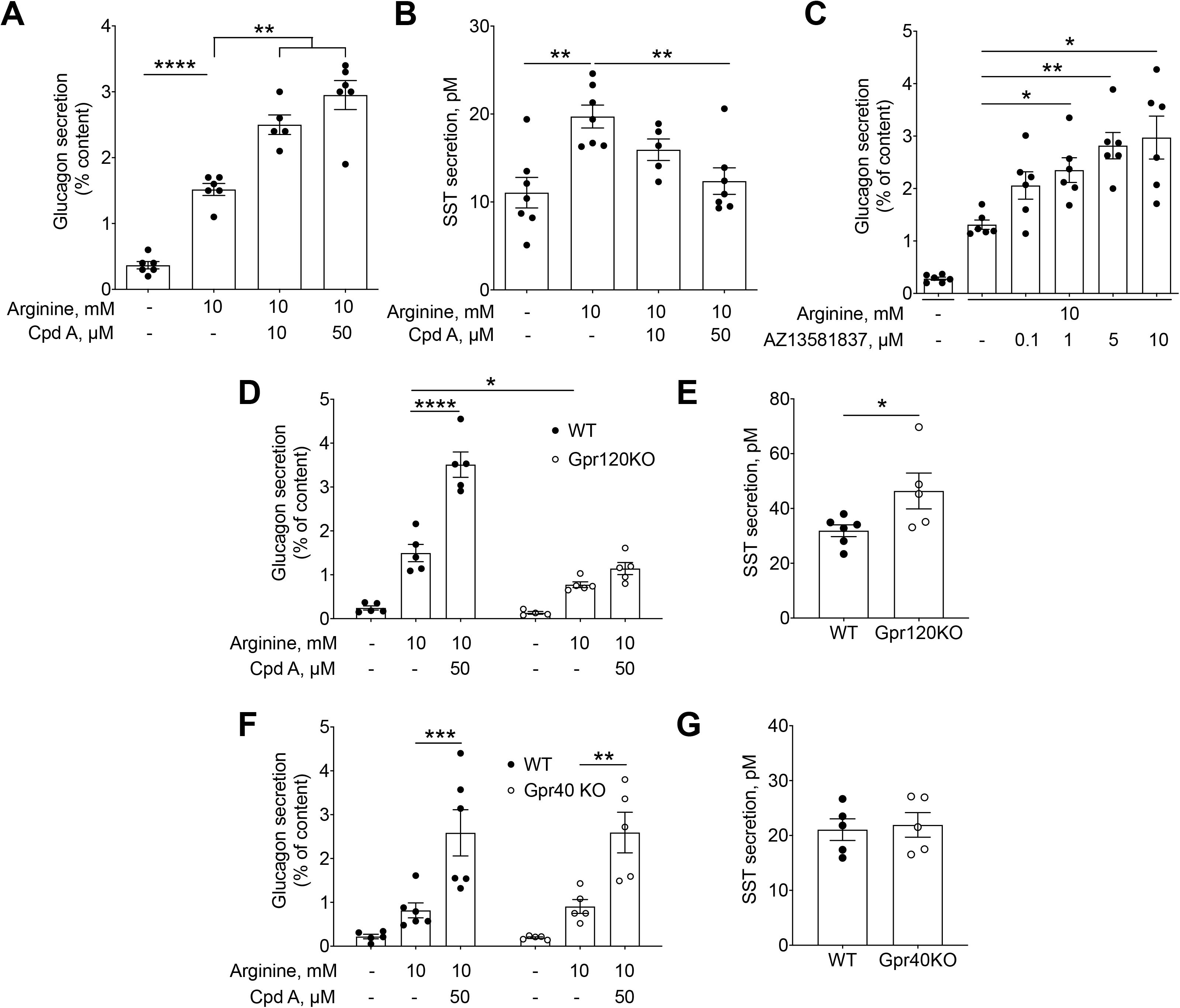
GPR120 activation potentiates arginine-stimulated glucagon secretion and inhibits SST secretion. Glucagon secretion presented as a percentage of islet glucagon content (A, C) and SST secretion measured in parallel on the same isolated islets (B) were assessed in 1-h static incubations in response to 10 mM arginine with or without the GPR120 agonists Cpd A (10-50 μM) (A, B) and AZ (0.1-10 μM) (C). Glucagon secretion in response to 10 mM arginine with or without Cpd A (50 μM) (D, F) and arginine-stimulated SST secretion in the absence of exogenous GPR120 agonist (E, G) was measured on isolated islets from Gpr120KO (D, E) and Gpr40KO (F, G) mice and WT littermate controls. Data represent individual values and are expressed as mean +/− SEM from 4-8 independent experiments. * P<0.05, ** P<0.005 or ***P<0.0005 between groups following one-way ANOVA (A-C) with Welch/Brown-Forsythe correction when necessary to compensate for SD variances, or two-way ANOVA (D-H) with Dunnett’s or Tukey’s post hoc adjustment for multiple comparisons.

### 3.4 *Gpr120* mRNA is enriched in δ cells in the mouse pancreas

To determine which islet cell types express *gpr120* we performed RNA in situ hybridization on sections of adult mouse pancreas. Double-fluorescence labelling with probes to *insulin*, *glucagon* and *sst* RNA in conjunction with the *gpr120* RNA probe was used to confirm *gpr120*-expressing cell identity. *Gpr120* RNA transcripts were detected predominantly in δ **(Fig. 4A & B)** and α cells **(Fig. 4C & D)**, with comparatively few in β cells (**Fig. 4E & F**). These observations are consistent with the expression pattern of FACS-sorted α, β and δ cells [31] by transcriptomic analysis (**Fig. 4G**) and with previous studies [28–30] showing that in the adult mouse pancreas *gpr120* expression is enriched in islet δ cells.

**Figure 4.**
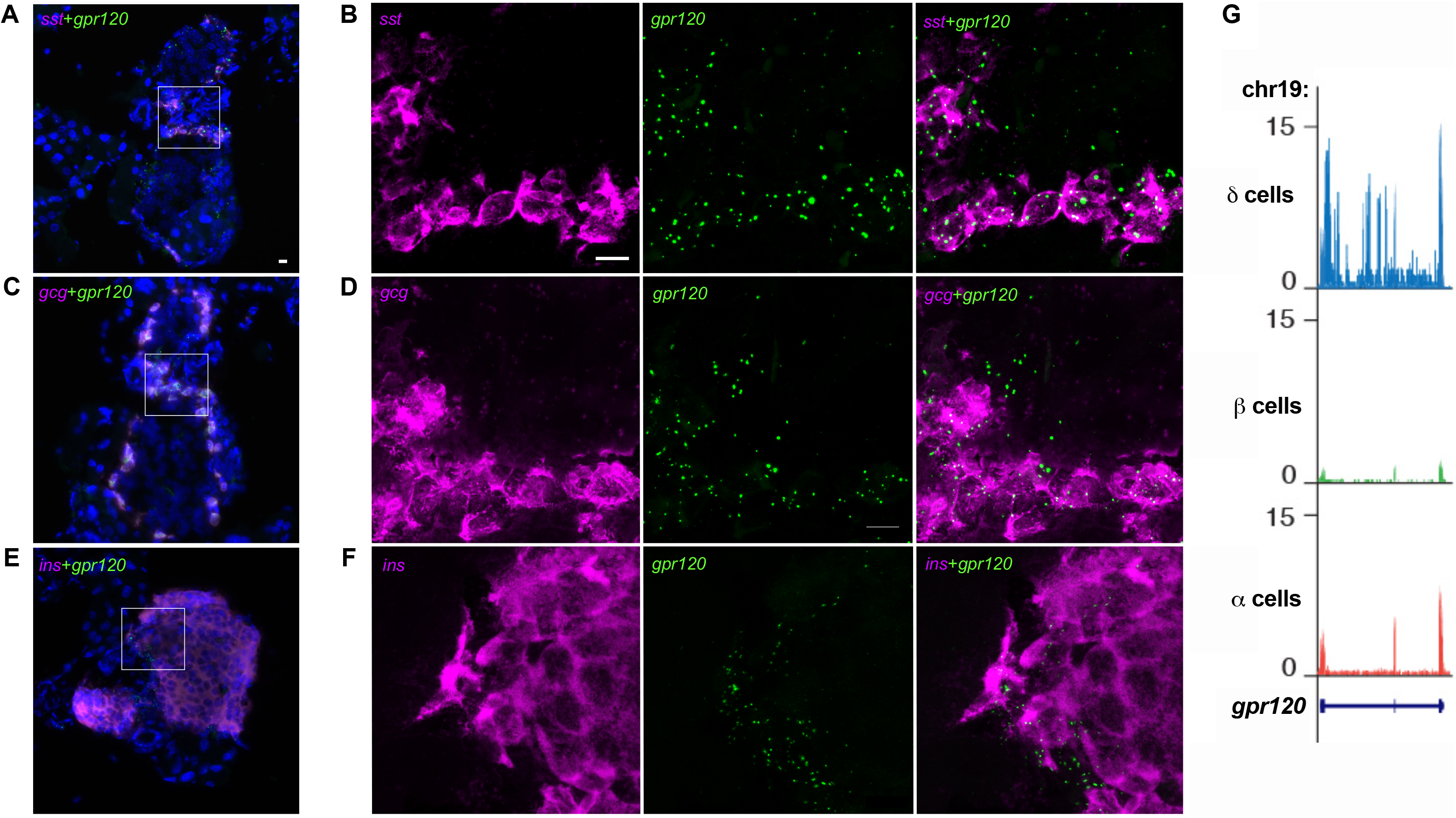
*Gpr120* mRNA is predominantly detected in mouse isletδ cells but also in someα and β cells. (A-F) *gpr120* (green) and *sst* (A, B), *glucagon* (*gcg*; C, D), or *insulin* (*ins*; E, F) (magenta) mRNA were detected in adult mouse pancreatic cryosections by fluorescent in situ hybridization. Representative sections are shown at 20 X (A, C, E) and 63 X (B, D, F) magnification. DAPI (blue). Scale bar = 10 μm. (G) Normalized browser plot illustrating *gpr120* mRNA expression in adult mouse islet cells based on transcriptomic analyses of FACS-sorted α, β and δ cells from reporter lines described in [31].

### 3.5 GPR120 activation potentiates cAMP generation and calcium fluxes in α and β cells but inhibits these signals in δ cells

To investigate signaling downstream of GPR120 we examined the effects of Cpd A on cAMP levels in α, β and δ cells in intact mouse islets using a cAMP reporter (CAMPER) expressed under the control of the glucagon, urocortin 3 (Ucn3) or SST promoter, respectively. Cpd A increased cAMP levels at 10 and 50 μM in the presence of 5.5 mM glucose in both α (**Fig. 5A & Supplementary Video 1**) and β (**Fig. 5B & Supplementary Video 2**) cells, albeit to a much lower level than the positive control forskolin. cAMP generation was also elevated in α cells in response to Cpd A in the presence of 5.5 mM glucose and arginine (**Supplementary Fig. 4 & Video 3**). In contrast, elevated cAMP levels in response to forskolin in the presence of 16.8 mM glucose were reduced upon addition of Cpd A in δ cells (**Fig. 5C & Supplementary Video 4**). As calcium is mobilized in response to GPR120 agonists in various cell types [6; 15; 22–24; 27; 38], we also measured calcium fluxes in α, β and δ cells in intact mouse islets using a calcium reporter (GCaMP6) expressed under the control of the glucagon, Ucn3 or SST promoter, respectively. In α cells, Cpd A and AZ augmented calcium fluxes, albeit mildly, induced by an amino acid mixture in the presence of basal (5.5 mM) glucose (**Fig. 6A**). The same α cells mounted robust calcium responses to AVP and KCl, as previously reported [37]. Calcium signals in the presence of 16.8 mM glucose were weakly increased by Cpd A in β cells (**Fig. 6B**). In contrast, Cpd A decreased calcium signals in δ cells in the presence of 5.5 mM glucose (**Fig. 6C**). Both β and δ cells exhibited a strong response to KCl confirming their viability. Taken together, these findings demonstrate that activation of GPR120 in islets leads to opposite effects in δ versus α and β cells. Whereas cAMP and calcium fluxes are decreased in δ cells, these signals are increased in α and β cells, fully consistent with the hormone secretion profile in response to GPR120 agonists.

**Figure 5.**
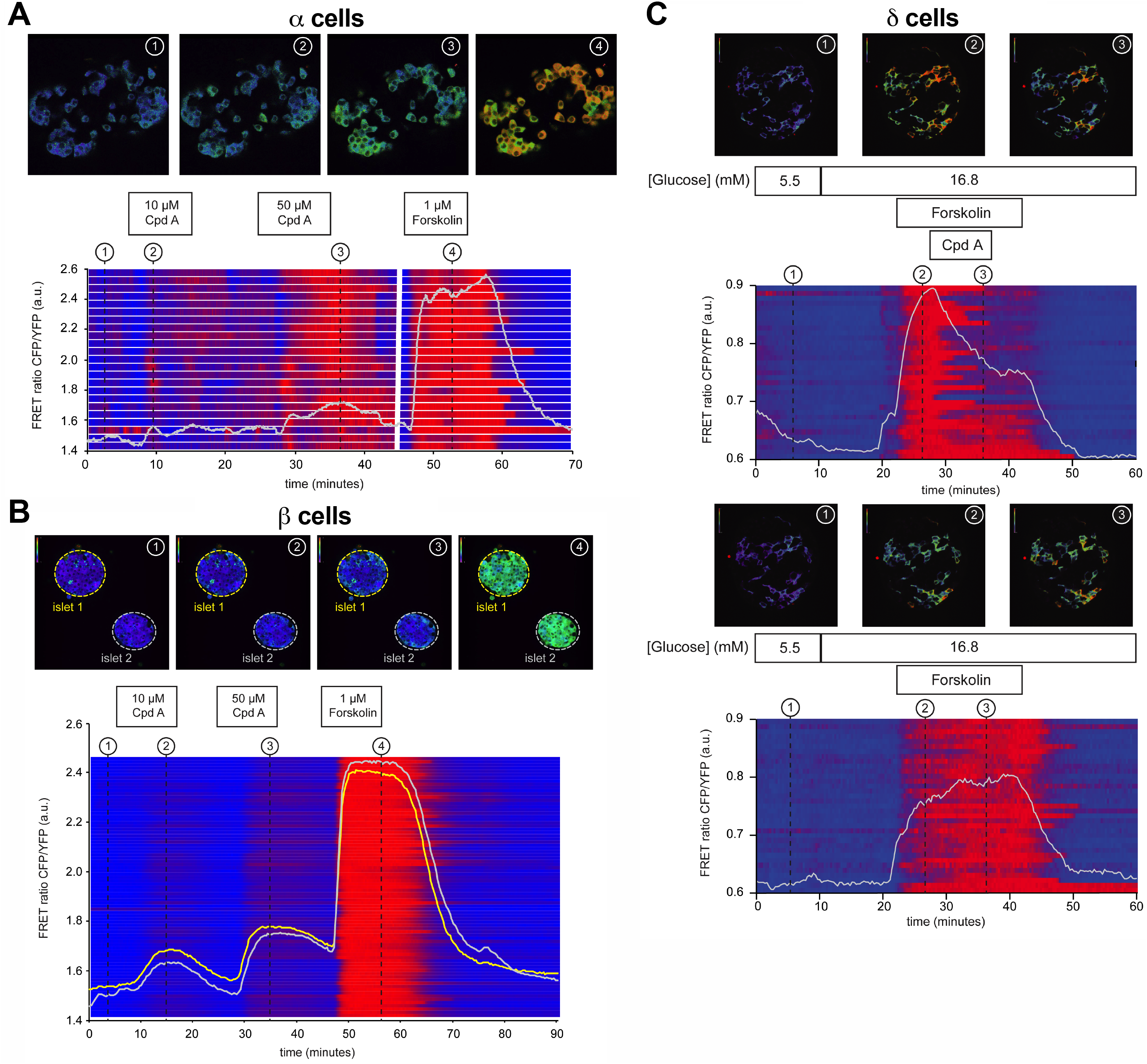
Cpd A increases cAMP levels inα and β cells but inhibits forskolin-induced cAMP elevation in δ cells. The cAMP sensor cAMPER was used to measure cAMP levels in individual α (A), β (B) and δ (C) cells in mouse islets. Cells were exposed to 5.5 mM glucose (A, B) or 5.5 mM followed by 16.8 mM glucose (C) and Cpd A at 10 μM (A, B) or 50 μM (A-C) and forskolin at 1 μM (A-C) at the times indicated.

**Figure 6.**
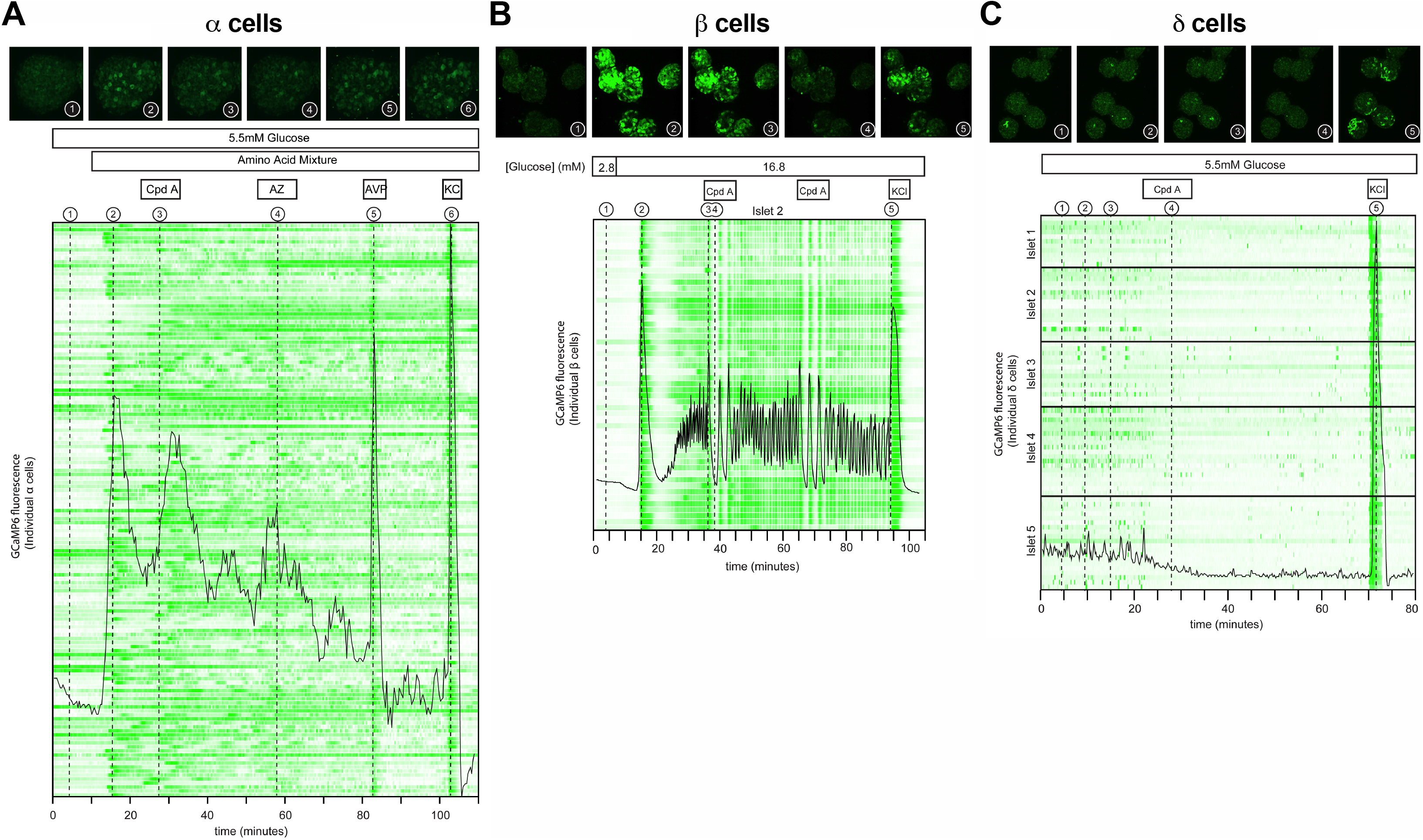
GPR120 agonists increase calcium signals in α and β cells but inhibit calcium signals and δ cells. The calcium sensor GCaMP6 was used to measure calcium activity in individual α(A), β(B) and δ(C) cells in the presence of 5.5 mM glucose and an amino acid mixture (2 mM each of glutamine, arginine and alanine) (A) or 2.8 mM glucose followed by 16.8 mM glucose (B) or 5.5 mM glucose (C). Cpd A (50 μM) and AZ (50 μM) were added at the times indicated. KCl (30 mM) induced depolarization (A-C) and AVP (10 nM) (A) served as positive controls.

### 3.6 GPR120 signaling in δ cells mediates the insulinotropic and glucagonotropic effects of Cpd A in isolated mouse islets

To determine the specific contribution of δ cell GPR120 signaling in the insulinotropic and glucagonotropic effect of Cpd A, we measured hormone secretion in static incubations in response to Cpd A in isolated islets from δGpr120KO mice compared to 3 control groups (WT, Gpr120Flox and SST-Cre). Cpd A at 50 μM significantly increased GSIS in WT, Gpr120Flox and SST-Cre islets, but not in δGpr120KO islets (**Fig. 7A)**. Accordingly, Cpd A was unable to inhibit GSSS in δGpr120KO islets (**Fig. 7B**). Cpd A increased arginine-induced glucagon secretion in WT, Gpr120Flox and SST-Cre islets, but not in δGpr120KO islets (**Fig. 7C)**. Cpd A significantly inhibited arginine-induced SST secretion in WT and Gpr120Flox islets but not in δGpr120KO islets (**Fig. 7D**). We were unable to determine the effect of Cpd A in SST-Cre islets as arginine-induced SST secretion was largely absent (**Fig. 7D**). Taken together, these results show that δ cell GPR120 signaling mediates the insulinotropic and glucagonotropic effects of Cpd A.

**Figure 7.**
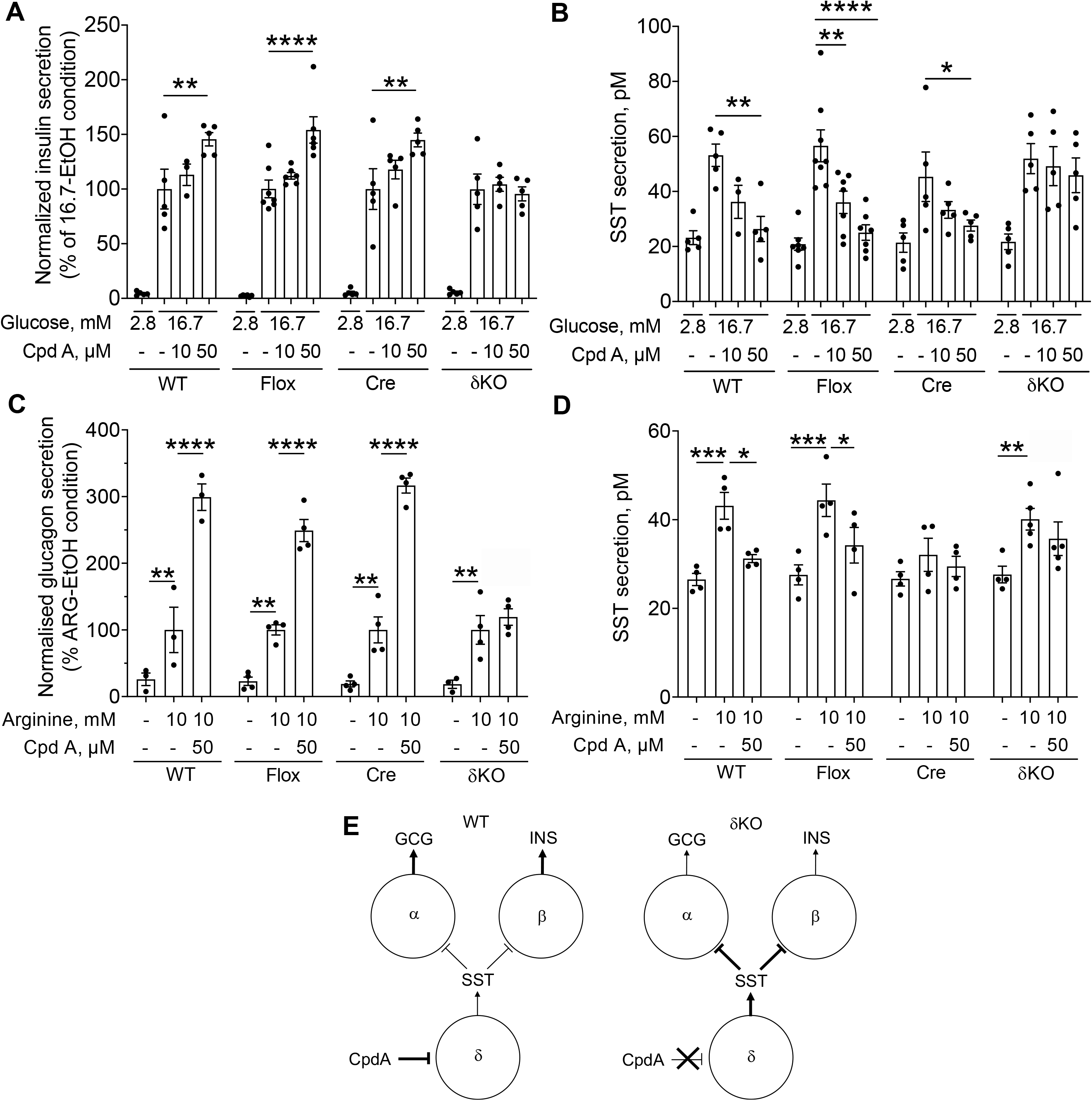
GPR120 signaling in δ cells mediates Cpd A-dependent potentiation of GSIS, arginine-stimulated glucagon secretion and inhibition of SST secretion. Normalized insulin secretion presented as a percentage of the 16.7-EtOH condition (A) and SST secretion measured in parallel on the same isolated islets (B) were assessed in 1-h static incubations in response to 2.8 or 16.7 mM glucose with or without Cpd A (10 or 50 μM) on isolated islets from WT, Gpr120^fl/fl^ (Flox), SST-Cre (Cre) and SST-Cre, Gpr120^fl/fl^ (δKO) mice. Normalised glucagon secretion presented as a percentage of the ARG-EtOH condition (C) and SST secretion measured in parallel on the same isolated islets (D) were assessed in 1-h static incubations in response to 10 mM arginine with or without Cpd A (50 μM) on isolated islets from mice of the 4 genotypes. Data represent individual values and are expressed as mean +/− SEM from 3-8 independent experiments. * P<0.05, ** P<0.005, ***P<0.0005 or ****P<0.0001 between groups following two-way ANOVA with Dunnett’s post hoc adjustment for multiple comparisons. (E) Web diagram illustrating the relationship between insulin, glucagon and SST secretion in WT and δGpr120KO (δKO) islets.

### 3.7 Inhibition of SST secretion by Cpd A is insensitive to PTX

As GPR120 is known to couple to inhibitory G proteins Gαi/o [10; 13], we asked whether the decrease in SST secretion in response to Cpd A can be reversed by pre-treating islets with PTX, an inhibitor of Gαi/o activity (**Fig. 8A**). The basal and glucose-induced increase in SST secretion were elevated in islets pretreated with PTX, consistent with Gαi/o inactivation alleviating tonic negative feedback from SST. However, the Cpd A-mediated repression of GSSS was unaffected by PTX exposure.

**Figure 8.**
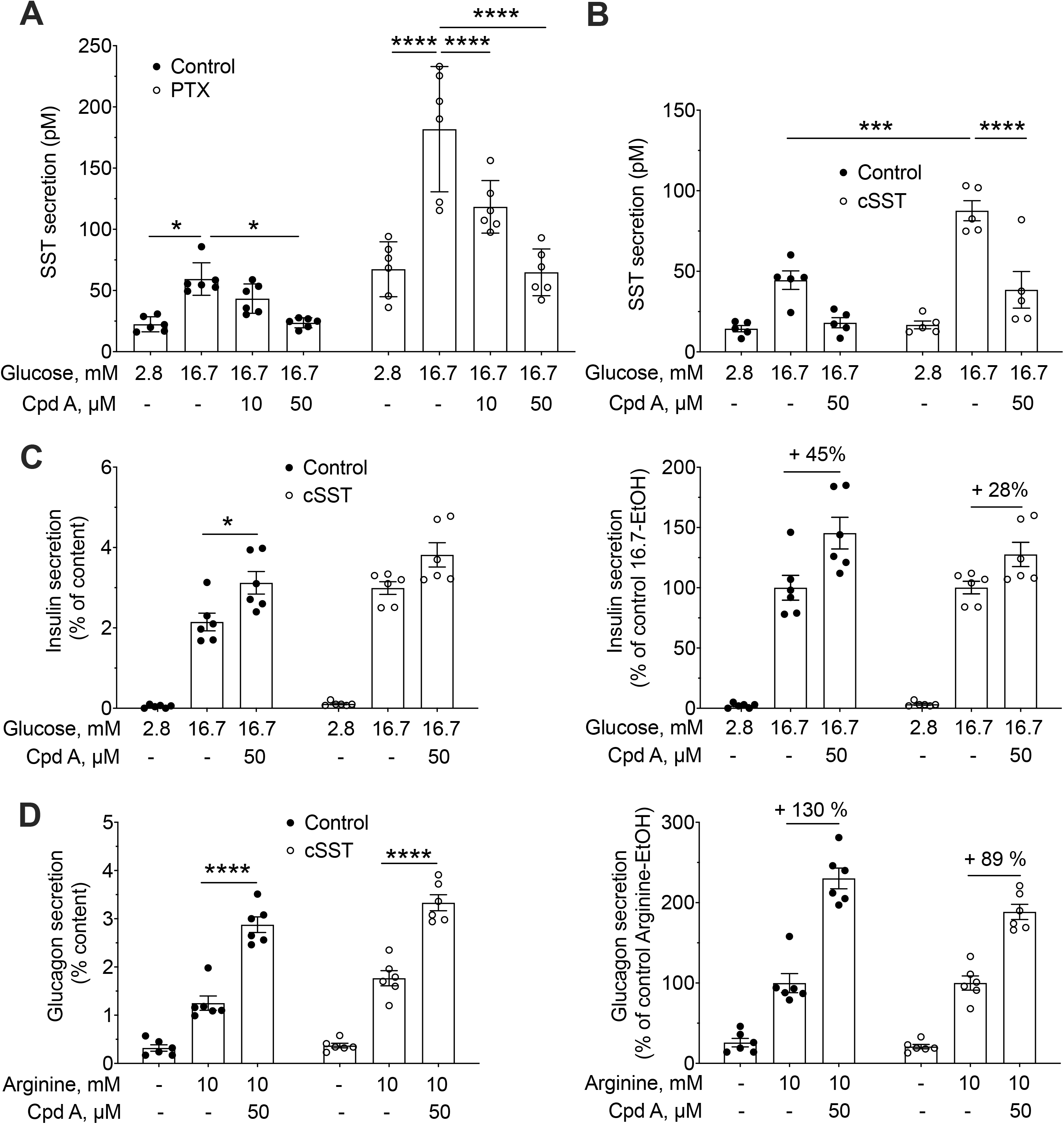
Cpd A-dependent inhibition of GSSS is not sensitive to PTX and potentiation of GSIS and arginine-stimulated glucagon secretion are only partially affected by SST receptor blockade. (A) SST secretion was assessed in 1-h static incubations in response to 2.8 or 16.7 mM glucose with or without the GPR120 agonist Cpd A (10 or 50 μM) following a 16-h pre-treatment in the presence or absence of PTX (100 ng/ml). (B-D) Effects of the SST receptor antagonist cSST (10 μM) on SST (B), insulin (C) and glucagon (D) secretion. (B) SST secretion was assessed in 1-h static incubations in response to 2.8 or 16.7 mM glucose with or without the GPR120 agonists Cpd A (50 μM) and/or cSST. (C) Insulin secretion presented as a percentage of islet insulin content was assessed as in (B). (D) Glucagon secretion presented as a percentage of islet glucagon content was assessed in 1-h static incubations in response to 10 mM arginine with or without the GPR120 agonists Cpd A (50 μM) and/or cSST. Data represent individual values and are expressed as mean +/− SEM of 5-6 independent experiments. * P<0.05, *** P<0.0005 or ****P<0.0001 between groups following two-way ANOVA with Dunnett’s or Tukey’s post hoc adjustment for multiple comparisons between all groups.

### 3.8 The regulation of islet hormone secretion by Cpd A is partially preserved upon SST receptor blockade

The stimulation of insulin and glucagon secretion by Cpd A in static incubations is inversely correlated with SST release (**Figs. 2, 3 & 7**) and is eliminated upon deletion of GPR120 in δ cells (**Fig. 7A, C & E**). Given the known paracrine, inhibitory effect of SST on insulin and glucagon secretion [39], our observations suggest that GPR120 activation alleviates SST inhibition of insulin and glucagon secretion. To directly assess this possibility, we tested the effects of Cpd A on hormone secretion in the presence of the competitive antagonist of all SST receptor isoforms, cyclosomatostatin (cSST). A large increase in GSSS was detected in the presence of cSST (**Fig. 8B**) suggesting that the SST negative feedback loop was alleviated. In contrast, the potentiation of GSIS (**Fig. 8C**) and arginine-stimulated glucagon secretion (**Fig. 8D**) by Cpd A was only partially eliminated in the presence of cSST. Although we did not directly confirm the effectiveness of cSST treatment, these data could suggest that the increase in insulin and glucagon secretion in response to CpdA involves a GPR120-mediated, δ cell-derived signal in addition to SST.

## 4. DISCUSSION

The objectives of this study were to clarify the role of islet GPR120 in the regulation of islet hormone secretion, and to delineate the specific contribution of δ-cell GPR120 signaling. We first showed that GPR120 activation improves glucose tolerance and increases insulin and glucagon secretion in vivo in mice. Accordingly, we found that GPR120 agonists inhibit SST secretion in isolated mouse islets and concomitantly potentiate GSIS and arginine-stimulated glucagon secretion. Importantly, the activity of the GPR120 agonist, Cpd A, is dependent on islet GPR120 but not GPR40. We then demonstrated that *gpr120* is enriched in δ cells and that GPR120 activation has opposing effects in δ versus α and β cells: whereas Cpd A reduced forskolin-induced cAMP generation and spontaneous calcium fluxes in δ cells, Cpd A increased both cAMP and calcium in α and β cells. Unexpectedly, inhibition of SST secretion by Cpd A was insensitive to PTX. Finally, we showed that the insulinotropic and glucagonotropic effects of Cpd A are lost in δ-cell specific *gpr120* KO islets and reduced by inhibition of SST receptor signaling. Overall, this study supports a predominant contribution of GPR120 signaling inhibiting δ-cells to relieve their inhibitory actions over insulin and glucagon secretion.

We found that acute oral administration of Cpd A increases glucose tolerance and insulin and glucagon secretion (**Fig. 1**). These data are in agreement with the study by Sundstrom et al. [23] demonstrating improved glucose tolerance and increased insulin secretion following administration of the GPR120 specific agonists AZ and Metabolex 36 in mice. Interestingly, Sundstrom et al. [23] demonstrated that the glucose lowering and insulinotropic effect of orally administered AZ and Metabolex 36 is dependent on gut-derived GLP-1. Hence, although our study supports a role of islet GPR120 in the control of insulin and glucagon secretion in response to Cpd A, enteroendocrine hormone secretion may also contribute to the in vivo effects of GPR120 agonism. Unfortunately, the specific contribution of GPR120 signaling in islet δ cells in vivo could not be assessed in our δGpr120KO mouse model as *gpr120* is deleted in all SST-expressing cells of the body. Conversely, GPR120-dependent GLP-1 release in vivo would not be expected to contribute positively to the glucagonotropic effects of Cpd A as GLP-1 has been shown to inhibit glucagon secretion from α cells [40].

In isolated islets from WT mice, the synthetic GPR120 agonists Cpd A and AZ, and the endogenous agonists ALA, EPA and DHA dose-dependently potentiate GSIS and inhibit GSSS. Furthermore, the effects of Cpd A are lost in *gpr120* but not *gpr40* KO islets (**Fig. 2**). These findings are in line with previous studies showing an insulinotropic effect of different GPR120 agonists in vitro in isolated rodent islets and insulin-secreting cell lines (INS-1E and BRIN-BD11) [22; 24–26]. In contrast Oh et al. [4] did not report a significant effect of Cpd A on GSIS in isolated mouse islets. However, the maximum dose tested was 10 μM, whereas in our hands a significant insulinotropic effect was detected beginning at 20 μM (**Fig. 2**). Similarly, Stone et al. [28] did not observe an increase in insulin secretion in response to the GPR120 agonist Metabolex 36 despite an inhibitory effect on GSSS. Our observation that low concentrations of Cpd A inhibit GSSS but do not significantly stimulate GSIS might explain why Stone et al. observed a reduction in GSSS without a concomitant increase in GSIS at their chosen concentration of Metabolex 36.

Deletion of GPR120 increases glucose- and arginine-stimulated SST secretion (**Fig. 2 & 3**). Although a constitutive, ligand-independent, activity of GPR120 could potentially account for these observations, we favor a role of endogenous, islet-derived, GPR120 ligands in controlling δ-cell function. This possibility would be analogous to the paracrine/autocrine action of long-chain saturated and monounsaturated FA such as palmitic, stearic, and oleic acids [41] and the arachidonic acid metabolite 20-HETE [42] which are released by the β cell and contribute to GPR40-mediated potentiation of GSIS. Whether these β cell-derived fatty acids also activate GPR120 in the δ cell will require further studies. Contrary to whole-body Gpr120KO islets, δGpr120KO islets did not secrete more SST in response to glucose compared to WT islets (**Fig. 7**). This may be explained by the lower levels of SST expression in several tissues of SST-Cre mice, due to the Cre knock-in allele behaving as a functional null allele [43]. Another possibility is that residual GPR120 activity in δ cells of δGpr120KO islets might contribute to the partial repression of SST secretion through the action of islet-derived GPR120 ligands.

Cpd A and AZ dose-dependently potentiate arginine-stimulated glucagon secretion (**Fig. 3**). Furthermore, the effect of Cpd A is dependent on GPR120 but not GPR40. Although Suckow et al. [27] also reported an increase in glucagon secretion in response to palmitate and DHA, the glucagonotropic effect in their study required both GPR120 and GPR40. The difference between our results and those of Suckow et al. is likely due to the activity of the GPR120 ligands at the concentrations tested. Hence, activation of GPR120 alone can be sufficient to promote glucagon secretion, as supported by recent studies using selective GPR120 agonists [26]. In the absence of exogenous GPR120 agonists, arginine-stimulated glucagon secretion is reduced in *gpr120*-but not *gpr40*-deficient islets pointing to a possible role of endogenous GPR120 ligands in the control of glucagon secretion.

We showed that the insulinotropic and glucagonotropic effects of Cpd A are mainly driven by δ-cell GPR120 signaling, as they are not observed in δGpr120KO islets (**Fig. 7)**. The enriched expression of *gpr120* mRNA in δ cells detected in pancreatic islet sections and in FACS-sorted islet cells [31] described herein (**Fig. 4**) and in previous *gpr120* mRNA expression studies [21; 28–30; 32] is consistent with these findings. Indeed, GPR120 activation in intact islets led to a decrease in forskolin-induced cAMP production and spontaneous calcium fluxes in δ cells both calcium and cAMP were increased in α and β cells (**Fig. 5 & 6 & Supplementary Fig. 4**). These observations are in full agreement with our secretion data, where Cpd A inhibited SST and potentiated nutrient-stimulated insulin and glucagon secretion. Collectively, these findings point to a model where GPR120-dependent inhibitory signals in δ cells suppress SST secretion which alleviates the paracrine inhibitory effects of SST on cAMP generation and calcium fluxes in both α and β cells. Nevertheless, *gpr120* mRNA and protein were also detected in α cells, and some β cells suggesting a possible direct regulation of glucagon and insulin secretion by GPR120. Mechanistically, any GPR120 expressed by α or β cells is clearly coupled to distinct downstream canonical signaling pathways compared to the inhibition of calcium and cAMP that characterizes GPR120 activation in δ cells. Furthermore, recent studies suggest that GPR120 is localized to primary cilia in mouse and human islet endocrine cells and that disrupting the cilial transport of GPR120 precludes agonist-dependent potentiation of insulin and glucagon secretion [26]. This finding is perfectly compatible with a role for GPR120 in δ cells in the regulation of insulin and glucagon secretion in islets. δ cells are also ciliated [26], and disruption of ciliary transport in islets would be expected to interfere with the GPR120-mediated inhibition of SST secretion. In addition, depletion of primary cilia in the β cell prevents SST inhibition of GSIS [44]. However, GPR120 agonists increase ciliary cAMP levels in clonal cell lines [26] an observation compatible with the increase in cAMP we report here in primary α and β cells. Hence, although Grp120 activation in α and β cells may contribute to glucagon and insulin secretion, our data strongly suggest that the predominant mechanism is δ-cell dependent.

Inhibition of SST secretion upon GPR120 agonism in δ cells (**Fig. 7**) suggests coupling to inhibitory G proteins, a possibility supported by the inhibition of forskolin-induced cAMP production and calcium mobilization in δ cells in response to Cpd A (**Fig. 5 & 6**). Surprisingly, inhibition of SST secretion by CpdA was completely insensitive to PTX (**Fig. 8**). This suggests that GPR120 might couple to PTX-insensitive inhibitory G proteins, such as Gαz, that is expressed alongside Gαi/o in δ cells [31] and also inhibits cAMP accumulation [45]. In contrast to our findings, Stone et al. [28] observed that inhibition of SST secretion in response to the GPR120 agonist Metabolex 36 in mouse islets was abrogated following PTX exposure suggesting the involvement of Gαi/o signaling. While this discrepancy may be due to biased agonism at GPR120 where Metabolex 36 and CpdA recruit divergent signaling pathways, further studies will be required to determine the identity of the inhibitory signals downstream of GPR120 in δ cells.

As GSSS is repressed in WT but not δGpr120KO islets, we inferred that Cpd A-induced insulin and glucagon secretion results mainly from the inhibition of SST secretion. However, the partial preservation of the stimulatory effect of Cpd A on hormone secretion in the presence of cSST (**Fig. 8**) leaves open the possibility that other, GPR120-dependent δ cell-derived signals may also play a role. Of note, the neuropeptide Y family gene peptide YY is expressed in δ cells [31; 46] and negatively regulates GSIS [46] and hence is a potential δ cell-derived signal that could partially mediate the effects of GPR120 activation on insulin secretion. Surprisingly, the effect of Cpd A on glucagon secretion is considerably stronger than its effect on insulin secretion, despite limited arginine-induced compared to glucose-induced SST secretion. Although, as discussed above, δ cell-derived signals other than SST might contribute to glucagon secretion, we favor the possibility that inhibition of SST secretion through activation of GPR120 in δ cells contributes to the glucagonotropic effect of Cpd A. Indeed, α cells are highly sensitive to δ cell-derived SST (SST14) due to expression of the SST receptor subtype, SSTR2 [47]. Furthermore, recent studies suggest that α-cell activity is intimately regulated by δ cells, which exert a tonic inhibition through SST [48–50].

Our data are consistent with the possibility that dietary and endogenous, including islet-derived, FA contribute to the regulation of glucagon and insulin levels in vivo by acting on GPR120 in δ cells. Interestingly, *gpr120* expression is reduced in islets from diabetic rodent models [24] and humans with type 2 diabetes [21]. Based on our findings, reduced δ cell GPR120 function would be expected to lead to elevated pancreatic SST levels, further limiting insulin and glucagon secretion. In dogs and rodents, pancreatic SST secretion is higher in diabetic compared to non-diabetic animals [51–53]. Furthermore, insulin levels and glycemic control are enhanced in β cell-specific SST receptor 5 KO mice [54] and following SSTR blockade [55]. Finally, SST receptor antagonists improve the glucagon counter-regulatory response to insulin-induced hypoglycaemia in diabetic rats [56; 57]. Although alterations in glucose control and islet hormone secretion were described in whole body Gpr120KO mice [3; 27; 58; 59], elucidating the role of δ cell GPR120 in glucose homeostasis will require in vivo analysis of δ cell-specific Gpr120KO mice which are not currently available.

In conclusion, our results show that the insulinotopic and glucagonotropic effects of islet GPR120 activation are largely mediated by inhibitory GPR120 signals in δ cell, in part but not exclusively through inhibition of SST secretion. These findings contribute to our understanding of the role of FA in the regulation of islet hormone secretion and of the mechanisms of action of putative anti-diabetic drugs targeting GPR120.

## 5. ACKNOWLEDGEMENTS

This work was supported by a Discovery Grant from the Natural Sciences and Engineering Research Council of Canada (RGPIN-2016-03952 to V.P.). M.O.H. received support from the National Institute of Diabetes and Digestive and Kidney Disease (NID DK-110276) and the Juvenile Diabetes Research Foundation (2-SRA-2019-700-S-B). M.L.C. was supported by a fellowship from the Société Francophone du Diabète and by a postdoctoral fellowship from the Montreal Diabetes Research Center. K.V. was supported by a postdoctoral fellowship from the Fond de Recherche Québec - Santé. G.M.N. was supported by a NIGMS-funded Pharmacology Training Program (T32GM099608). VP holds the Canada Research Chair in Diabetes and Pancreatic Beta-Cell Function.

We thank Maria Sörhede Winzell and Linda Sundström from Astra Zeneca for providing AZ-13581837; Jean Rivier from the Salk Institute for Biological Studies for synthesizing and providing SST and AVP; and Grace Fergusson, Mélanie Éthier, Annie Levert and Hasna Maachi from the CRCHUM for valuable technical assistance.

## 6. AUTHOR CONTRIBUTIONS

**Marine L. Croze**: Conceptualization, Methodology, Investigation, Formal Analysis, Writing – Original Draft; **Marcus Flisher**: Methodology, Investigation, Formal Analysis, Visualization; **Arthur Guillaume**: Investigation, Formal Analysis; **Caroline Tremblay**: Methodology, Investigation; **Glyn M. Noguchi**: Investigation; **Sabrina Granziera**: Investigation; **Kevin Vivot**: Methodology, Investigation; **Vincent C. Castillo**: Investigation; **Scott A. Cambpell**: Investigation; **Julien Ghislain**: Conceptualization, Validation, Writing – Review and Editing, Supervision; **Mark O. Huising**: Conceptualization, Validation, Writing – Review and Editing, Supervision, Funding Acquisition; **Vincent Poitout**: Conceptualization, Validation, Writing – Review and Editing, Supervision, Funding Acquisition, Project Administration.

## Footnotes

1 ALA: α-Linolenic acid; AVP: arginine vasopressin; AZ: AZ-13581837; cAMP: cyclic AMP; Cpd A: Compound A; cSST: cyclosomatostatin; DHA: docosahexaenoic acid; EPA: eicosapentaenoic acid; FA: fatty-acids; FACS: fluorescence-activated cell sorting; GSIS: glucose-stimulated insulin secretion; GSSS: glucose-stimulated somatostatin secretion; KCl: potassium chloride; KO: knock-out; PP: pancreatic polypeptide; PTX: pertussis toxin; SST: somatostatin; WT: wild-type

