## Supplementary Tables and Figures for "Free fatty-acid receptor 4 inhibitory signaling in delta cells regulates islet hormone secretion in mice"

**CROZE ET AL. - SUPPLEMENTARY MATERIAL LEGENDS**

**Supplementary Figure 1. *Gpr120* gene expression in Gpr120KO and δGpr120KO islets.** *gpr120* mRNA was measured in isolated islets from prev-Flox (A), Gpr120KO (C), δGpr120KO (δKO, D) and control mice. mRNA was determined by quantitative RT-PCR and normalized to *cyclophilin A.* Data are presented as the fold-change over WT and represent individual values expressed as mean ± SEM of 3-8 independent experiments. SST secretion in response to 2.8 or 16.7 mM glucose with or without Cpd A (10 and 50 µM) was measured on isolated islets from prev-Flox (B) mice and WT littermate controls. * $p<0.05$, ** $p<0.005$, ***$p<0.0005$ or ****$p<0.0001$ as compared to WT (A, C, D) or 16.7-EtOH condition (B).

**Supplementary Figure 2. Insulin secretion and content in isolated islets exposed to GPR120 agonists.** Insulin secretion (A, C, E, G, I) and content (B, D, F, H, J) were assessed in 1-h static incubations in response to 2.8 or 16.7 mM glucose with or without the Gpr120 agonists Cpd A (10-50 µM) (A, B, G-J), AZ (0.1-10 µM) (C, D) or the ω-3 fatty acids, alpha-linolenic acid (ALA), eicosapentaenoic acid (EPA) and docosahexaenoic acid (DHA) (100 µM) (E, F) in WT (A-J), Gpr120KO (G, H) and Gpr40KO (I, J) islets. Data represent individual values and are expressed as mean +/- SEM of 4-8 independent experiments. * $p<0.05$, ***$p<0.0005$ or ****$p<0.0001$ between groups following one-way ANOVA (A-F) or two-way ANOVA (G-J) with Dunnett's (A-F, versus 16.7-EtOH condition) or Tukey's (G-J) post hoc adjustment for multiple comparisons and Welch/Brown-Forsythe correction when necessary to compensate for SD variances.

**Supplementary Figure 3. Glucagon secretion and content in isolated islets exposed to GPR120 agonists.** Glucagon secretion (A, C, E, G) and content (B, D, F, H) were assessed in 1-h static incubations in response to 10 mM arginine with or without the GPR120 agonists Cpd A (10-50 µM) (A, B, E-H) or AZ (0.1-10 µM) (C, D) in WT (A-H), Gpr120KO (E, F) and Gpr40KO (G, H) islets. Data represent individual values and are expressed as mean +/- SEM from 4 to 8 independent experiments. * $p<0.05$, ** $p<0.005$ or **** $p<0.0001$ between groups following one-way ANOVA (A-D) or two-way ANOVA (E-H) with Dunnet's or Tukey's post hoc adjustment for multiple comparisons, and Welch/Brown-Forsythe correction when necessary to compensate for SD variances.

**Supplementary Figure 4. Cpd A potentiates cAMP levels in arginine treated α cells.** The cAMP sensor cAMPER was used to measure cAMP levels in individual α cells in mouse islets. Cells were exposed to arginine (10 mM) and Cpd A (50 µM) in the presence of 5.5 mM glucose at the times indicated.

**Supplementary Videos. Cpd A potentiates cAMP levels in α and β cells but inhibits forskolin-induced elevation in cAMP in δ cells.** The cAMP sensor cAMPER was used to measure cAMP levels in individual α (Video 1 and 3), β (Video 2) or δ (Video 4) cells following exposure to 5.5 mM glucose (Video 1 and 2), 5.5 mM glucose followed by 10 mM arginine (Video 3) or 2.8 mM followed by 16.8 mM glucose (Video 4) after which Cpd A (10 and 50 µM) and forskolin (1 µM) were added at the times indicated.

### Supplementary Figure 1

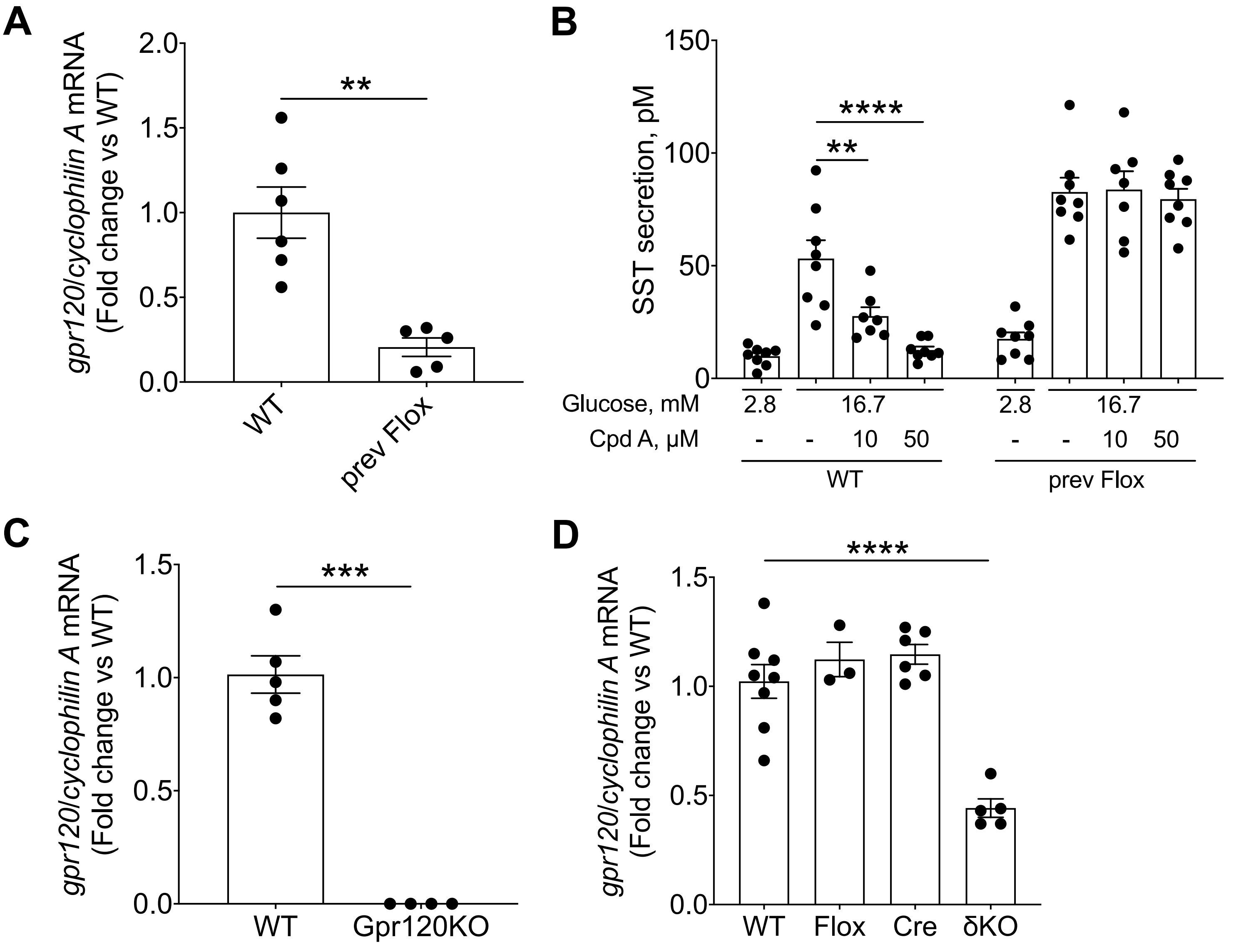

### Supplementary Figure 2

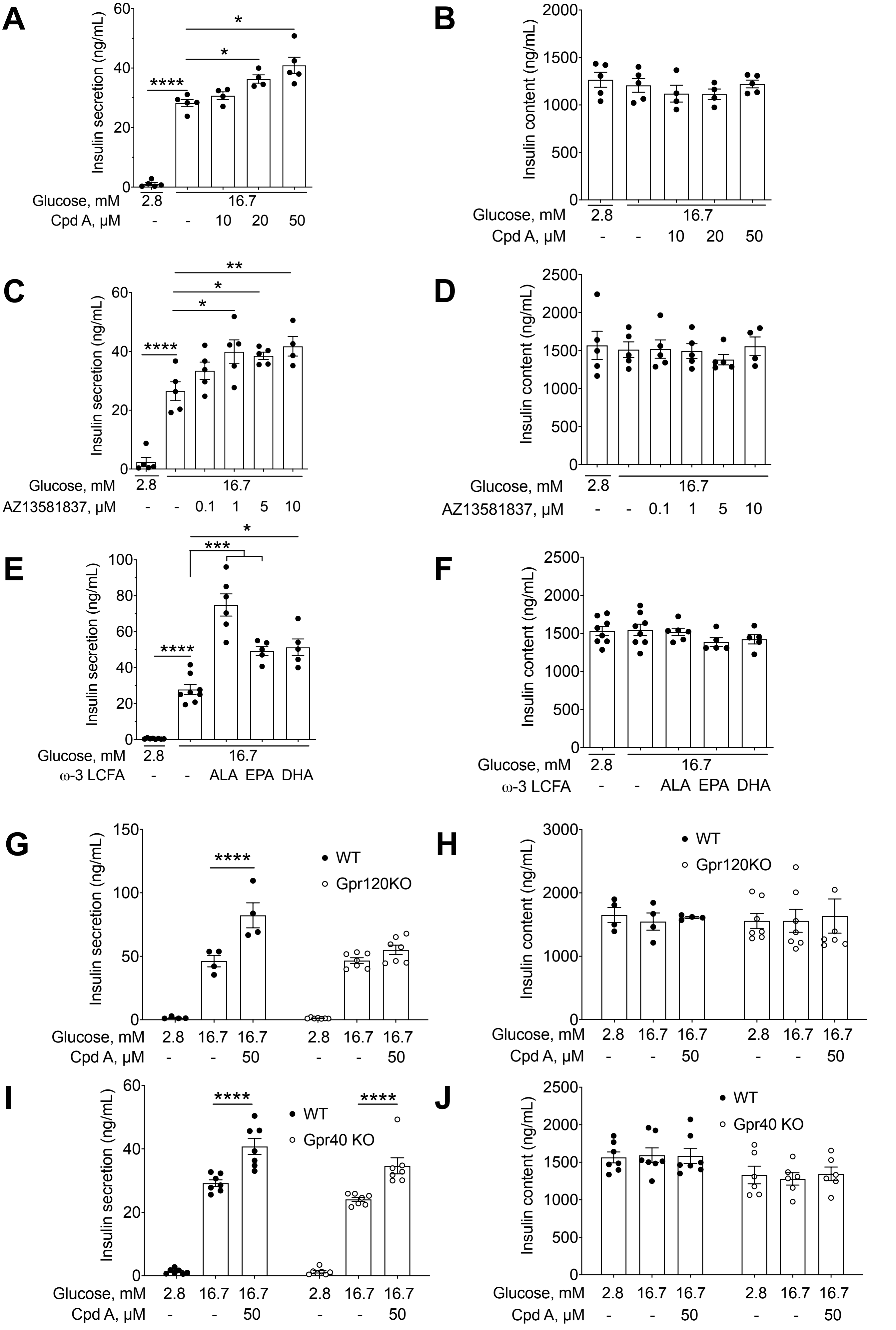

### Supplementary Figure 3

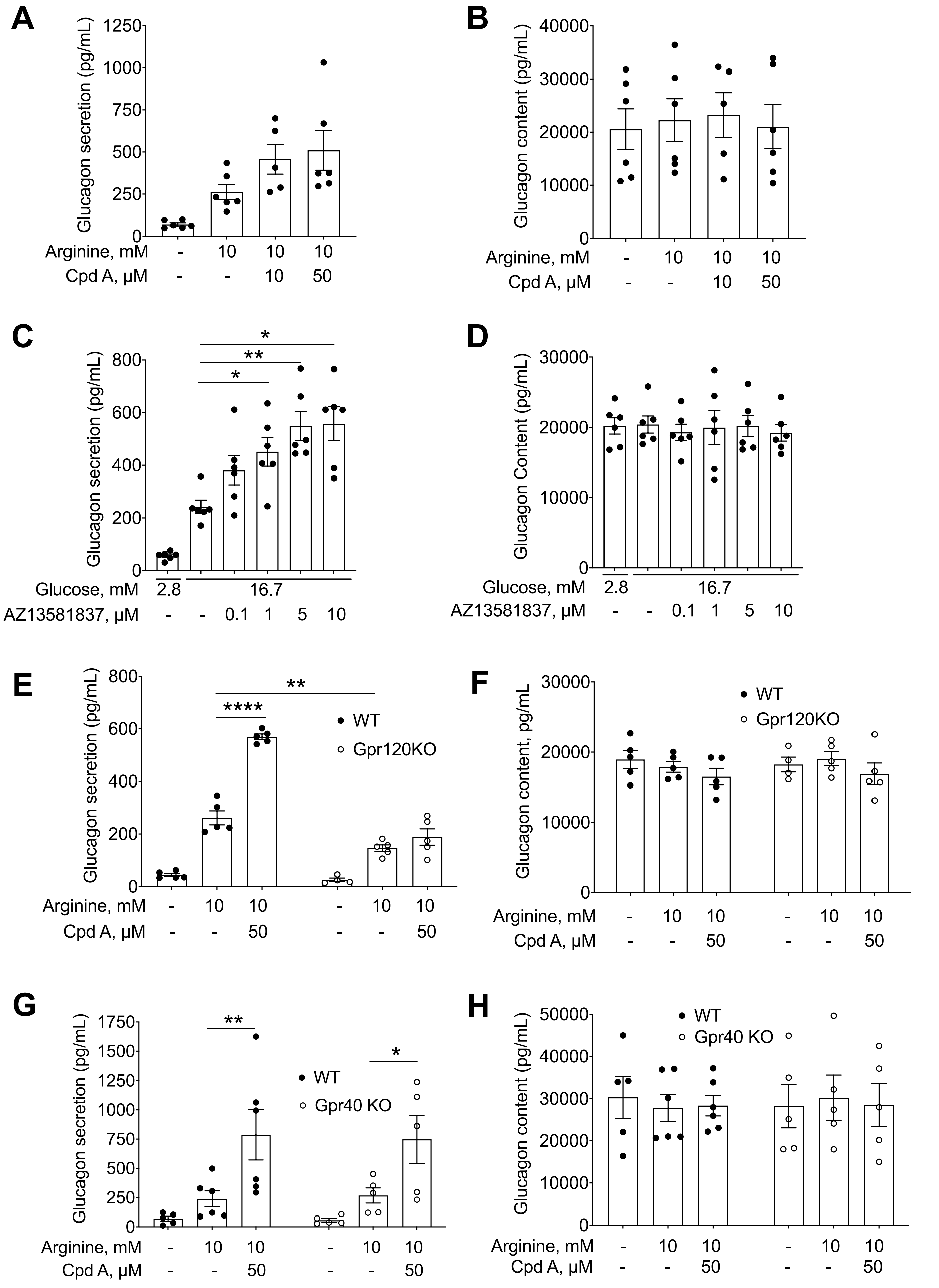

### Supplementary Figure 4

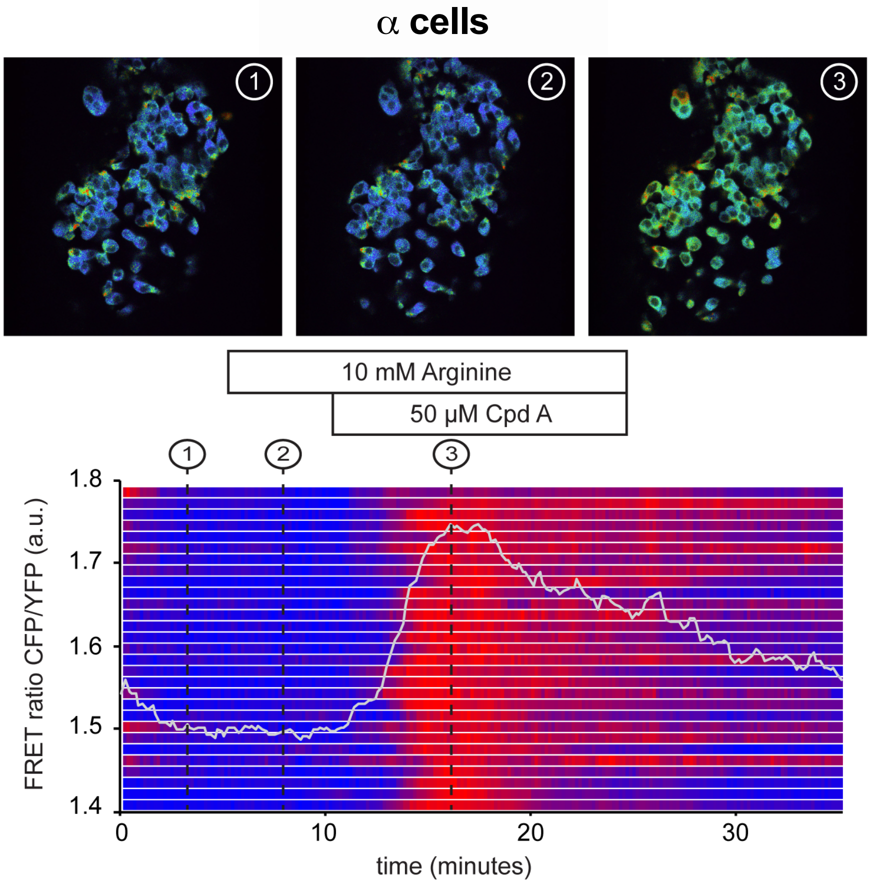

**Supplementary Table 1 - Primer sequences used for genotyping**

| | Primer 1 | Primer 2 | Primer 3 |
|---|---|---|---|
| **Gpr40** | GCAGCGCATCGCCTTCTATC | GGCTGGACAACAGTACCAGTTCC | ACGCTTGTCCTCTCCAGGTGG |
| **Gpr120D** | TCCTCGACGGTATCGATAAGCTA | GGCTGTAAGCTCCCATTGCT | |
| **Gpr120flox** | AAACTGCAGAGACTATTATGAGGC | ATCCCACTACTGTAGGATTTCTCC | |
| **SST-Cre** | TGGTTTGTCCAAACTCATCAA | TCTGAAAGACTTGCGTTTGG | |
| **Cre** | ATGTCCAATTTACTGACCG | CGCCGCATAACCAGTGAAAC | |
| **NNT** | GTGGAATTCCGCTGAGAGAACTCTT | GGGCATAGGAAGCAAATACCAAGTTG | GTAGGGCCAACTGTTTCTGCATGA |

**Supplementary Table 2 - Primer sequences used for real-time PCR**

| Gene Symbol | Gene Name | Forward Primer | Reverse Primer |
|---|---|---|---|
| Ffar4 | Free fatty acid receptor 4 | TGCCCCTCTGCATCTTGTTC | GGTTGGGCCAATCCAATGTG |
| Ppia | Peptidylprolyl isomerase A | TCTGAGCACTGGAGAGAAAGG | TTCTCTCCGTAGATGGACCTG |
